# Personalized stimulation therapies for disorders of consciousness: a computational approach to inducing healthy-like brain activity based on neural field theory

**DOI:** 10.1101/2024.05.20.594934

**Authors:** Daniel Polyakov, P.A. Robinson, Eli J. Müller, Glenn van der Lande, Pablo Nunez, Jitka Annen, Olivia Gosseries, Oren Shriki

## Abstract

Disorders of consciousness (DoC) pose significant challenges in neurology. Conventional neuromodulation therapies for DoC have exhibited limited success, with varying effectiveness among patients. In this study, we introduce a computational approach for constructing personalized stimulus signals capable of inducing healthy-like neural activity patterns in DoC patients.

Leveraging a simplified brain model based on neural field theory, we fit this model to the power spectrum of a patient with DoC and derive a personalized stimulus time series to induce a healthy-like power spectrum. By applying this stimulus to brain regions typically targeted by stimulation therapies such as deep brain stimulation and repetitive transcranial magnetic stimulation, we demonstrate *in silico* the ability of our method to elicit EEG power spectra resembling those of healthy individuals.

We speculate that in the course of a long-term treatment, when the brain produces healthy-like activity, it may trigger intrinsic plasticity mechanisms, potentially leading to sustained improvements in the patient’s condition. While further clinical adjustments and validation are needed, this novel approach offers promise in tailoring brain stimulation therapies for DoC patients. Moreover, it presents potential extensions to other conditions that could also benefit from brain stimulation therapies.

**Author summary:** In this research, we tackled a challenging issue in the field of neurology - disorders of consciousness (DoC). These conditions, which include states like coma and the minimally consciousness state, have been difficult to treat effectively. While traditional brain stimulation treatments have been tried, they do not always work well, and their effects can vary greatly from one person to another. Thus, we set out to find a new way to help these patients. Our approach involves using computer models of the brain to design personalized treatments. We designed a method to generate stimulation signals tailored to individual DoC patients. These signals aim to trigger healthier brain activity patterns, similar to those found in individuals without consciousness disorders. What is exciting is that if the brain begins to produce these healthy-like patterns, it could potentially lead to lasting improvements in the patient’s consciousness level. We show that this approach works in different simulations, and can be applied in multiple neuromodulation techniques like deep brain stimulation and transcranial magnetic stimulation. This research opens up new possibilities for more effective treatments for people dealing with these challenging disorders. Plus, it might work for other conditions treated with similar brain stimulation therapies.

## Introduction

### Disorders of consciousness

Disorders of consciousness (DoC) encompass a range of conditions characterized by impaired awareness and wakefulness resulting from severe brain damage. These conditions include coma, unresponsive wakefulness syndrome (UWS), and minimally conscious state (MCS). Collectively, these conditions disrupt the intricate neural processes that underlie human consciousness irrespective of arousal, challenging our fundamental understanding of how the brain forms awareness, and cognition, thereby prompting profound scientific, clinical, and ethical inquiries [1, 2].

DoC can originate from a variety of causes, such as traumatic brain injuries, anoxia, strokes, and intoxication. These conditions lead to impairments in a persons’ awareness and in their ability to respond to the external environment [1]. However, in clinical practice, the assessment of a patient’s level of consciousness is primarily based on their responsiveness. For DoC assessment of the level of consciousness, the Coma Recovery Scale-Revised (CRS-R) is typically used. It assesses auditory, visual, motor, oromotor, communication, and arousal functions. Each function contributes points to a total score ranging from 0 (coma) to 23 (healthy), reflecting overall severity. Diagnosis of specific DoC states relies on achieving specific response levels within each item category, such as visual pursuit or command response, or by using a CRS-R index based on the highest item in every category [3, 4].

The consciousness level of DoC patients can be enhanced through various therapeutic approaches. These approaches include neurorehabilitation strategies, surgical procedures, pharmacological interventions, brain stimulation, and emerging experimental treatments such as stem-cell therapy [5–7]. In this context, our focus is on brain stimulation, specifically the use of electrical or magnetic stimulation to modify brain activity, in order to promote neural plasticity and to re-activate damaged brain circuits [8].

### Brain Stimulation Therapies

Various stimulation therapies have been explored as potential interventions to improve outcomes in individuals with DoC, focusing particularly on those with UWS or MCS. Techniques such as deep brain stimulation (DBS), transcranial magnetic stimulation (TMS), transcranial direct current stimulation (tDCS), transcranial alternating current stimulation (tACS), and vagus nerve stimulation (VNS) have been applied to DoC patients to increase cortical excitability and, consequently, to enhance consciousness (see Fig. 1) [6, 8, 9].

**Fig 1.**
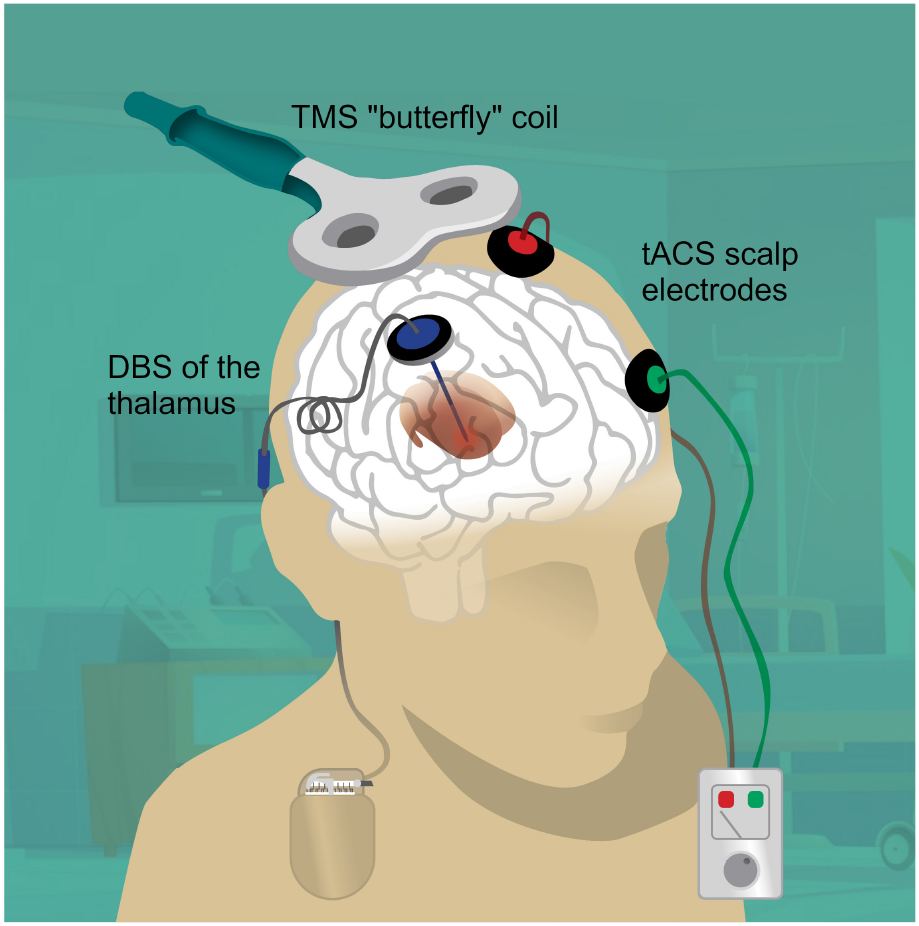
Example of various brain stimulation methods applied to a patient. DBS involves the implantation of electrodes into specific brain areas that deliver controlled electrical impulses. TMS utilizes magnetic pulses that are directed through the patient’s skull. tACS transmits alternating electrical current between two electrodes positioned on the patient’s scalp. (This figure includes a hospital background image by Upklyak from Freepik.)

Among invasive therapies, DBS requires surgically implanting electrodes into specific brain regions to deliver controlled electrical impulses, showing promise in improving consciousness among individuals with DoC. Stimulation of various thalamic nuclei has demonstrated significant success rates, with sustained improvements observed in numerous patients [10–14]. VNS therapy, involving stimulation of the vagus nerve, offers a less invasive alternative, potentially modulating consciousness by influencing corticothalamic networks and the reticular activation system via connections with the brainstem [15]. Despite limited spatial precision, VNS has shown a sustained improvement in the level of consciousness in MCS patients [8, 16].

TMS is a non-invasive technique that involves delivering magnetic fields through the skull to induce electrical currents in specific brain regions. Repetitive TMS (rTMS) [17] has demonstrated efficacy in enhancing behavioral scores among DoC patients. However, its efficacy varies among individuals, and long-term benefits remain uncertain [8, 9, 18]. In both tDCS and tACS, mild electrical current is transmitted through the scalp, with the former aiming to modify cortical excitability and the latter seeking to synchronize with brain oscillations. Evidence indicates that tDCS can lead to significant improvements in CRS-R scores, but like rTMS, the long-term efficacy and effects of tDCS are subjects of ongoing research [6, 8, 9]. As for tACS, there is not enough research to draw conclusions about its effectiveness for the treatment of DoC [19]. Notably, all of these non-invasive techniques typically target the dorsal prefrontal cortex, which is part of a network associated with consciousness in the brain [8, 18–20].

Despite the potential of these brain stimulation methods, their adoption as reliable therapies for DoC faces challenges due to the considerable variability in treatment outcomes, attributed to factors like the patient’s condition, the extent of brain damage, and individual biological characteristics [8, 14]. In turn, this leads to a large variability in the patients’ need in stimulation, from the most appropriate technique (e.g., transcranial, or more invasive DBS), to stimulation parameters in terms of e.g., location, frequency and duration. With the large heterogeneity and low sample sizes, finding a common, effective stimulation paradigm has major feasibility challenges. Computational modeling is an outstanding tool in overcoming these shortcomings, because without increased burden for the patient, a search for optimal stimulation could be conducted. To address this issue, we aim to provide a road to improvement in these issues by modeling an abstract representation of the discussed stimulation techniques and focusing on tailoring the electro-magnetic signal for specific patients. This stimulation approach is coupled to a physiologically-inspired brain model capable of representing the patient’s own brain activity. By tailoring the stimulation to each patient’s unique profile, we hope to demonstrate how modeling could be utilized in the future to deal with the DoC population’s heterogeneity that could affect treatment effectiveness.

### Multiscale brain modeling

Over the past half-century, significant progress has been made in the development of brain activity models that address various phenomena across different brain regions and spatial scales. In particular, efforts have been directed toward creating comprehensive multiscale models of whole-brain activity that are capable of simulating various phenomena and generating realistic signals. One noteworthy modeling framework, “The Virtual Brain”, employs neural mass models to represent brain regions interconnected based on connectome data. This model demonstrates the capacity to simulate diverse phenomena, including epilepsy and sleep stages. Nevertheless, fitting it to experimental data remains challenging because the use of non-linear differential equations is required to generate the activity of each region [21, 22]. Another approach, the “Multiscale Neural Model Inversion” framework, focuses on effective connectivity. It constructs interregional networks based on structural connectivity and fits their activity to functional magnetic resonance imaging data through functional connectivity correlation, ultimately generating blood-oxygen-level-dependent signals similar to those that are observed in pathologies like major depression and Alzheimer’s disease. However, the output aligns only with the first and second moments of the experimental data, thereby capturing only the fundamental properties of the simulated phenomena [23]. In contrast, neural field theory (NFT) modeling offers an efficient approach to building networks of interconnected brain regions that interact through physiologically-based wave equations, thereby capturing both stationary and transient neural activity. NFT has demonstrated success in representing consciousness-related phenomena like sleep stages and epilepsy and has presented substantial capabilities of classifying states of consciousness [24–27]. Furthermore, the NFT framework provides convenient tools for fitting and simulating neural activity using standard computing equipment [26, 28].

Moreover, it is important to note that NFT has previously been employed to model various stimulation techniques and their impacts on both healthy and pathological brains. For instance, Wilson et al. (2016, 2018) used NFT to model the effects of TMS on calcium-dependent synaptic plasticity in long-term potentiation and depression processes [29, 30]. Muller et al. (2018) applied NFT to model the effects of DBS for Parkinson’s disease, including a detailed model of the corticothalamic-basal ganglia system that replicated key clinical features of the disease, such as high beta-activity. They explored different DBS protocols and successfully demonstrated their ability to reduce beta-activity in the model [31, 32]. Muller et al. (2023) also investigated consciousness modulation through DBS by simulating the expression of propofol anesthesia in a corticothalamic system and introducing a stimulus to the matrix thalamus. Their model demonstrated arousal from anesthesia, characterized by wake-like dynamics that facilitated high information transfer [33]. Thus, NFT appears to be a suitable framework for developing personalized stimulation signals for DoC patients. Furthermore, it offers the capability to incorporate these signals into the modeled stimulation techniques mentioned here or those that may be modeled in the future.

### Proposed approach

As previously mentioned, our approach aims to establish a systematic method for constructing patient-specific and region-specific stimulation signals. The core idea involves utilizing a corticothalamic NFT model (CTM) capable of simulating both healthy-like and DoC-like neural activity, while allowing for targeted stimulation of the modeled regions. We fit this model to a specific DoC patient, and then we mathematically derive the stimulus signal required to induce typical healthy-like neural activity within the model. The fitting of the model and the construction of the stimulus signal rely on the corresponding EEG power spectra, aiming to generate a healthy-like power spectrum. We apply the constructed signal to brain regions typically targeted by DBS, rTMS, and tACS, and successfully demonstrate the effectiveness of our method *in silico*.

Our approach holds promise for application in DBS, rTMS, and tACS, as they demonstrated the ability to influence various bands of DoC patients’ power spectrum [11, 19, 34]. Conversely, tDCS is not well-suited for our approach because it only allows for amplitude adjustment, which is insufficient for independently controlling the power of different frequencies. Moreover, since the brainstem and vagus nerve are not included in the CTM, we did not construct a signal for application in VNS in this study. Generally, with appropriate adaptations, this method has the potential to significantly advance a wide range of brain stimulation therapies, offering a personalized approach that could lead to more effective treatments for DoC.

## Materials and methods

### Neural field theory

Neural-field modeling [24, 35, 36] stands as a tool for constructing physiologically inspired brain models capable of mimicking various multiscale measures of brain activity. This approach captures a continuum of corticothalamic activity by simulating the local dynamics in each population and employing wave equations to describe the propagation between these populations. [37]. Consequently, it effectively replicates and integrates numerous phenomena observed in EEG data, including spectral peaks observed during waking and sleeping states, event-related potentials, measures of connectivity, and spatiotemporal structure, and even the dynamics of epileptic seizures (see Fig. 2) [25, 38–44]. The model’s parameters encompass various biophysically meaningful quantities such as synaptic strengths, excitatory and inhibitory gains, propagation delays, synaptic and dendritic time constants, and axonal ranges. NFT represents a bottom-up approach to whole-brain modeling, which involves averaging over microstructure to derive mean-field equations. This method effectively complements analyses conducted at the cellular and local neural network levels. Additionally, NFT’s strength lies in its dynamic analysis, which explores the model’s states and their stability in relation to changes in model parameters [26, 42, 43].

**Fig 2.**
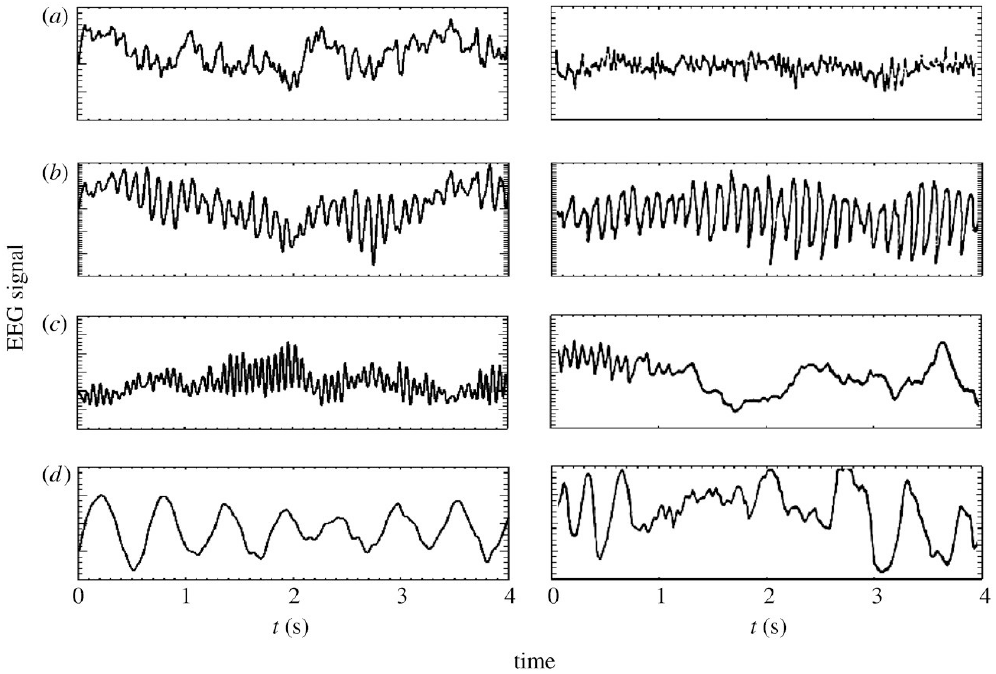
Simulated vs. experimental EEG time series of waking and sleeping states. In the left panels: model generated time series of (a) eyes-open resting state, (b) eyes-closed resting state, (c) sleep-stage 2, and (d) sleep-stage 3. In the right panels: corresponding time series from human subjects [45, 46]. (The figure was originally published by Robinson et al. (2005) [24].)

The neural field *ϕ*(**r**, *t*) [s^*−*1^] represents a spatiotemporal neural activity propagating among neural populations when averaged across scales of approximately 0.1 millimeters. Within the framework of the CTM, four distinct neural populations are involved, with key connectivities, illustrated schematically in Fig. 3. These populations comprise excitatory (*e*) and inhibitory (*i*) cortical neurons, thalamic relay nuclei neurons (*s*), thalamic reticular nucleus neurons (*r*), and sensory inputs (*n*). Each of these populations has a soma potential *V*_*a*_(**r**, *t*) [V] that is influenced by contributions *ϕ*_*b*_ from presynaptic populations, and generates outgoing neural activity *ϕ*_*a*_(**r**, *t*).

**Fig 3.**
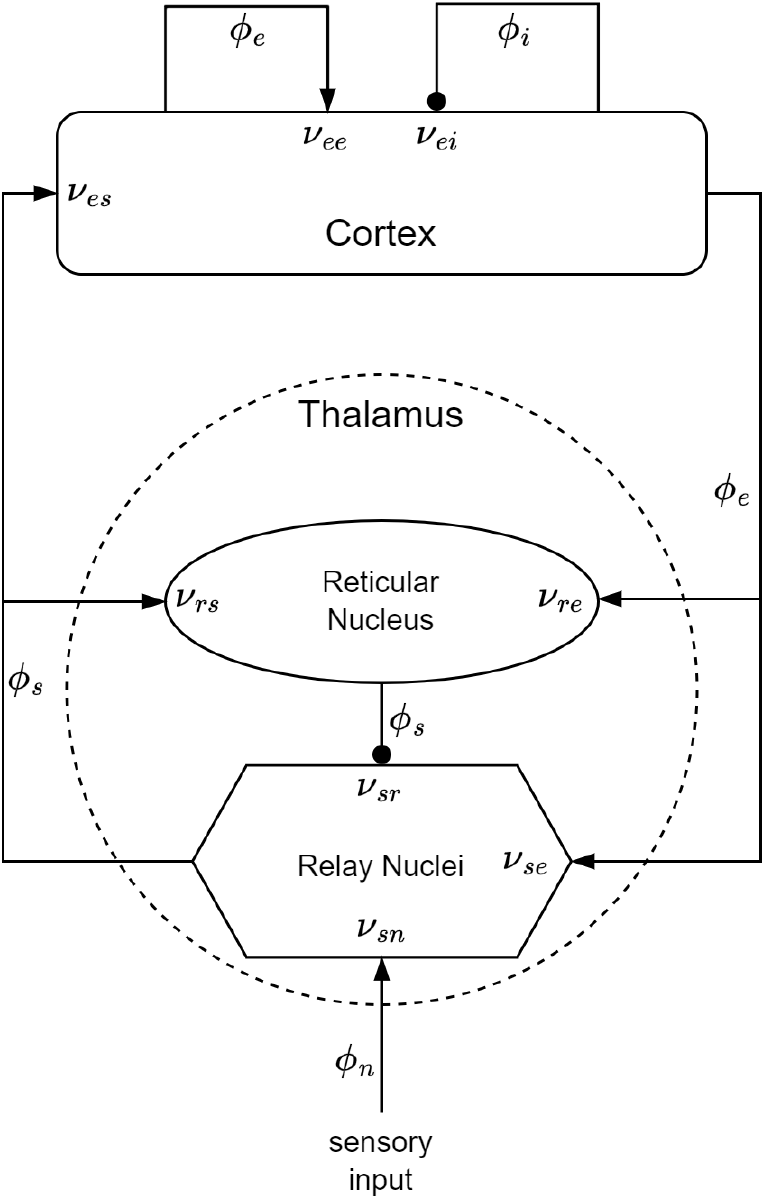
The CTM diagram. The neural populations shown are cortical excitatory, *e*, and inhibitory, *i*, thalamic reticular nucleus, *r*, and thalamic relay nuclei, *s*. The parameter *ν*_*ab*_ quantifies the strength of the connection from population *b* to population *a*. Excitatory connections are indicated by pointed arrowheads, while inhibitory connections are denoted by round arrowheads.

The dendritic spatiotemporal potential *V*_*ab*_ [V] is linked to the input *ϕ*_*b*_ through Eq. 1. The parameter *ν*_*ab*_ = *s*_*ab*_*N*_*ab*_ [V *·* s] represents the strength of the connection from populations *b* to *a*, where *N*_*ab*_ is the mean number of synapses per neuron *a* from neurons of type *b*, and *s*_*ab*_ [V *·* s] is the mean time-integrated strength of soma response per incoming spike. The parameter *τ*_*ab*_ [s] refers to the one-way corticothalamic time delay, and *D*_*a*_(*t*) is a differential operator, as described in Eq. 2. Here, 1*/α* and 1*/β* denote the characteristic decay time and rise time, respectively, of the soma response within the corticothalamic system.

Dendritic component:

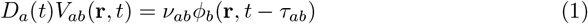

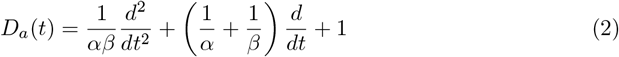

The soma potential *V*_*a*_ is determined as the sum of its dendrite potentials, as outlined in Eq. 3. This potential undergoes some smoothing effects attributed to synaptodendritic dynamics and soma capacitance. Furthermore, the population generates spikes at a mean firing rate *Q*_*a*_ [s^*−*1^], which is related to the soma potential through a sigmoid function *S*(*V*_*a*_) (relative to the resting state), as shown in Eq. 4. In this equation, *Q*_max_ denotes the maximum firing rate, while *θ* and 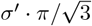 correspond to the mean and the standard deviation (SD), respectively, of the firing threshold voltage.

Soma component:

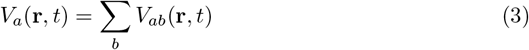

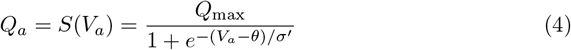

The field *ϕ*_*a*_ approximately follows a damped wave equation with a source term *Q*_*a*_, as detailed in Eq. 5. The differential operator *D*_*a*_(**r**, *t*) is defined in Eq. 6, where *v*_*a*_ [m *·* s^*−*1^] represents the propagation velocity, *r*_*a*_ [m] denotes the mean range, and *γ*_*a*_ = *v*_*a*_*/r*_*a*_ [s^*−*1^] signifies the damping rate.

Axonal component:

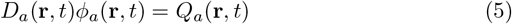

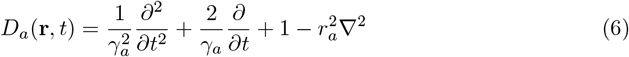

In this model, only *r*_*e*_ is large enough to induce notable propagation effects.

Consequently, the fields of other populations can be approximated as *ϕ*_*a*_(**r**, *t*) = *S*[*V*_*a*_(**r**, *t*)]. Additionally, we assume that the only non-zero time delays between populations are *τ*_*es*_, *τ*_*is*_, *τ*_*se*_, and *τ*_*re*_= *t*_0_*/*2, where *t*_*0*_ is the total time it takes to traverse the corticothalamic loop. It is important to note that Eq. 5 encompasses the corticocortical time delays, as the wave equation inherently accounts for delays arising from propagation across the cortex. To further simplify the model, we assume random intracortical connectivity, leading to *N*_*ib*_ = *N*_*eb*_ for all *b* [47]. This assumption implies that the connection strengths are also symmetric, resulting in *ν*_*ee*_ = *ν*_*ie*_, *ν*_*ei*_ = *ν*_*ii*_, and *ν*_*es*_ = *ν*_*is*_ [24, 26]. *Numerical integration [28] or, when feasible, analytical integration [48] of NFT equations produces a spatiotemporal activity signal that propagates across the cortical surface*.

### CTM EEG power spectrum

In scenarios of spatially uniform steady-state activity, it is possible to analytically compute the power spectrum of the model, eliminating the necessity for numerical integration. The steady state is attained by setting all time and space derivatives to zero. Employing the first term of the Taylor expansion enables a linear approximation of all potential perturbations from the steady state. Applying a Fourier transform to the model equations under these conditions yields Eq. 7 for the dendritic component and Eq. 8 for the axonal component. Within these equations, *ω* = 2*πf* denotes the angular frequency, *k* = 2*π/λ* signifies the wave vector (*λ* is the wavelength), and 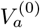 represents the steady-state potential [26].

Dendritic equations in the frequency domain:

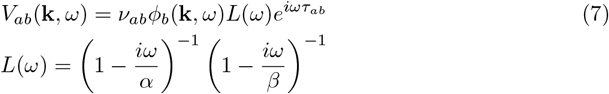

Axonal equations in the frequency domain:

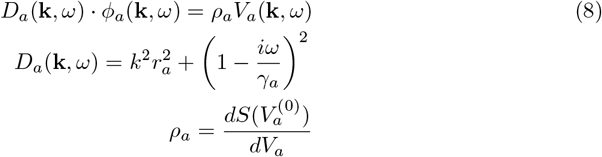

Using Eq. 3, we can write Eq. 8 as:

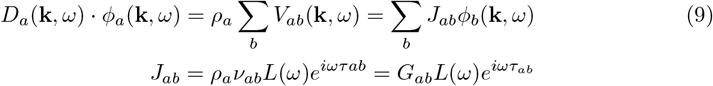

Then we can represent the interactions among the different populations within the CTM in matrix form:

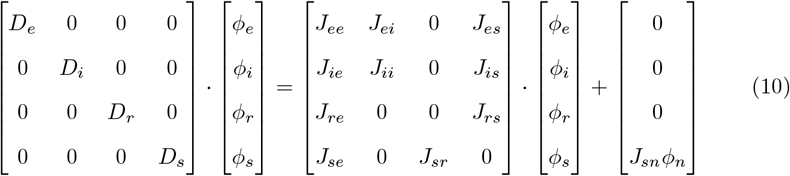

Eq. 10 can also be written in a compact form, when **J**^**⋆**^***ϕ***^**⋆**^ is the external input to the CTM:

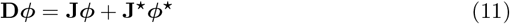

By solving Eq. 10, considering all the previously mentioned assumptions regarding *D*_*a*_, *ν*_*ab*_, and *τ*_*ab*_, we can derive Eq. 12. In this context, the quantities *G*_*ese*_ = *G*_*es*_*G*_*se*_, *G*_*esre*_ = *G*_*es*_*G*_*sr*_*G*_*re*_, and *G*_*srs*_ = *G*_*sr*_*G*_*rs*_ correspond to the overall gains for the excitatory corticothalamic, inhibitory corticothalamic, and intrathalamic loops, respectively. The firing rate of sensory inputs to the thalamus, *ϕ*_*n*_, is approximated by white noise. Without loss of generality, *ϕ*_*n*_(*ω*) can be set to 1, while only *G*_*sn*_ is subject to variation.

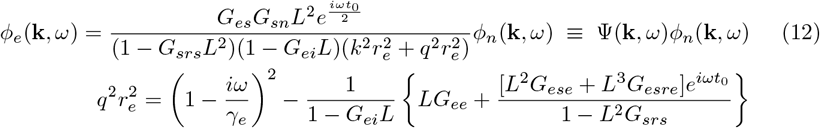

The excitatory field *ϕ*_*e*_ is considered a good approximation of scalp EEG signals [26]. The EEG power spectrum *P* (*ω*) (Eq. 13) is calculated by integration of |*ϕ*_*e*_(**k**, *ω*)|^2^ over **k** when the cortex is approximated as a rectangular sheet of size *L*_*x*_ *× L*_*y*_. When considering periodic boundary conditions, this integral transitions into a summation over spatial modes with a discrete *k*. The filter function *F* (*k*) serves as an approximation of the low-pass spatial filtering that occurs due to volume conduction through the cerebrospinal fluid, skull, and scalp.

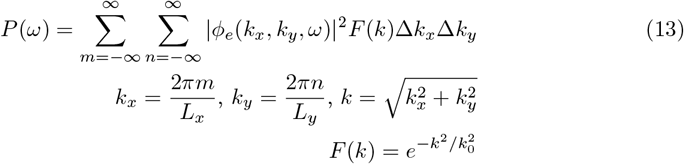

By merging Eq. 13 with Eq. 12, we can formulate the expression for the power spectrum of the CTM. Given that *ϕ*_*n*_ is equal for every *k*, we can write it as *ϕ*_*n*_(*ω*) outside the summation.

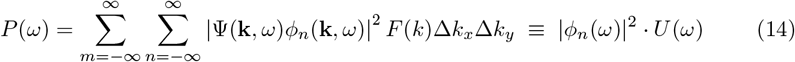

### Model fitting and signal generation

The analytical form of the CTM power spectrum, as expressed in Eq. 14, can be fitted to the experimental EEG power spectrum. This fitting process is carried out through a Monte-Carlo Markov-Chain optimization procedure applied to model parameters, a method implemented in the *braintrak* toolbox [26]. Here, we optimize the following model parameters: *G*_*ee*_, *G*_*ei*_, *G*_*es*_, *G*_*se*_, *G*_*sr*_, *G*_*sn*_, *G*_*re*_, *G*_*rs*_, *α, β, t*_*0*_, and *EMG*_*a*_. To support multi-electrode fitting, we assumed that *G*_*ee*_, *G*_*ei*_, *G*_*sn*_, *α,β*, and *t*_*0*_ exhibit a cosine-like variation over the scalp [37]. Subsequently, time series were generated from the fitted model by numerically integrating its partial differential equations over time, incorporating the *NFTsim* toolbox [28]. An illustrative example showing an experimental EEG Cz electrode power spectrum, a power spectrum derived from the fitted model, and a simulated EEG Cz electrode power spectrum is depicted in Fig. 4. The Cz electrode was specifically chosen to illustrate spectra of simulated data throughout this work, as its activity effectively represents the overall EEG activity generated by the CTM.

**Fig 4.**
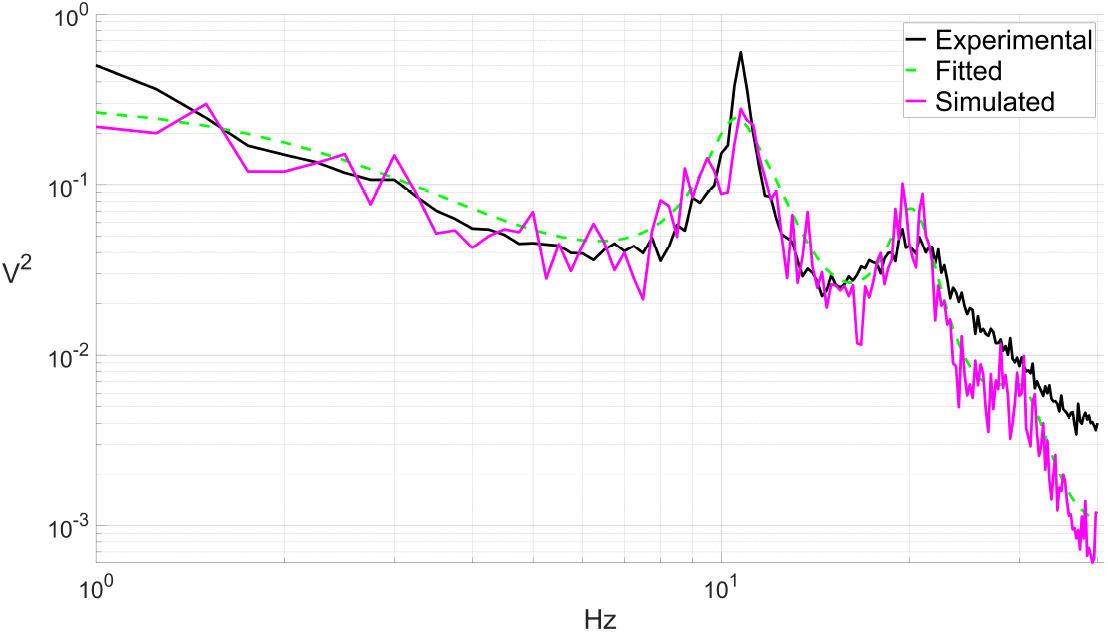
Experimental, fitted, and simulated power spectra. Example of a Cz electrode EEG power spectrum of a healthy subject (black curve), a fitted CTM power spectrum (green dashed curve), and a Cz electrode power spectrum of a simulated EEG (purple curve).

### Stimulus construction workflow

We recruited NFT equations and a framework to derive a stimulus signal that would cause a CTM fitted to a specific DoC patient to produce a power spectrum similar to that of a healthy individual. Here is an outline of our workflow:

1. Compute the power spectra of a typical healthy subject and a specific DoC patient.
2. Fit CTMs to both the healthy subject and DoC patient, and obtain the corresponding power spectra, *P* ^*hlt*^(*ω*) and *P* ^*doc*^(*ω*).
3. Compute the stimulus time series by solving *P* ^*hlt*^(*ω*) = *P* ^*doc*+*stim*^(*ω*). Present the time series as a Fourier series to support continuous stimulation.
4. Generate an EEG time series from “Healthy-fitted” and “DoC-fitted with stimulation” models, and compare 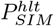 to 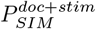.

It is worth noting that we can also choose to skip the fitting of the healthy subjects’s data and use the experimental power spectrum 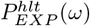 directly, as the stimulus-specific equations do not rely on the healthy-fitted CTM. We present both approaches in our results, providing flexibility in the experimental design.

### Experimental data

In our research, we utilized a dataset consisting of open-eyes resting-state EEG recordings acquired from DoC patients in various states, as well as from healthy controls. Each participant’s EEG activity was recorded using a Hydro-Cel GSN high-density electrode net (Electric Geodesics, EGI) equipped with 256 electrodes, with a sampling rate of 250Hz. Data collection took place in a controlled environment, characterized by darkness and minimal sensory stimuli, with EEG recording sessions lasting approximately 30 minutes. Behavioral assessments using the CRS-R were performed several times within a few days, including just before the EEG acquisition [49].

We chose an MCS patient (a 29-year-old male with a history of traumatic brain injury, CRS-R score of 9, and 181 months since the injury) and a UWS patient (a 31-year-old female with a history of brain hemorrhage, CRS-R score of 6, and 45 months since the injury) along with two healthy subjects (#1: a 62-year-old male; #2: a 19-year-old male) to present the application of our method. It is important to emphasize that these subjects were chosen for illustrative purposes only, and our approach is applicable to any pair of subjects, as shown further.

Each subject’s EEG data went through several processing steps. First, we kept only 137 central electrodes and removed the rest in order to focus only on scalp-based electrodes. The signals from these electrodes were high-pass filtered above 0.2Hz and notched at 50Hz to eliminate line noise. Subsequently, to streamline computational processes, they were down-sampled to 125Hz. The 30-minute recording was then partitioned into nearly stationary sections (required for CTM fitting, which is performed through steady-state activity power spectrum), with durations varying from 2.5 to 10 minutes. Detecting potential onsets of stationary sections involved splitting the electrode signals into 10-second segments and identifying when the segment’s SD deviated by two SDs compared to the preceding segment. Defining stationary section boundaries required simultaneous onsets in more than 5% of the electrodes, with at least 2.5 minutes between onsets. Sections lasting between 10 and 20 minutes were equally divided into two, while those between 20 and 30 minutes were split into three sections.

The cleaning process for each stationary section of the data involved several sequential steps. Initially, noisy channels (electrode signals) were identified by segmenting the data into 5-second segments and calculating the SD of each channel within a segment. A channel within a segment was flagged as noisy if its SD exceeded seven times the SD of the entire non-segmented channel, or if it was three times higher than that of the other channels within the same segment. Segments with at least one noisy channel were deemed noisy, and entire channels were marked as noisy if more than 33% of their segments exhibited noise. These identified entire noisy channels were then removed and replaced with channels generated through a spherical spatial interpolation of the remaining channels. Subsequently, the data was re-referenced to the average of all channels and any remaining artifacts were eliminated. Artifacts were defined as 1-second segments where any sample’s SD was five times greater than the channel’s entire SD. To further refine the data, we conducted an independent component analysis to isolate and remove components associated with eye movements and non-EEG activity. After decomposition, we employed the *IClabel* toolbox [50] to retain only those components identified with at least 50% confidence as originating from genuine neural activity. The steps involving spatial interpolation, average referencing, and independent component analysis were all executed using the *EEGLAB* toolbox [51].

From the various stationary sections of each subject, we retained sections where fewer than 10% of the channels were flagged as noisy and fewer than 10% of the independent components were removed during the cleaning process. Among these sections, we selected the one with the least amount of detected artifacts per minute of recording. Subsequently, we computed multi-electrode power spectra for the selected stationary section using a fast Fourier transform. Then, we fitted the power spectrum in the frequency range of 1–40Hz to a CTM, as described above.

### Personalized stimulus derivation

We can easily integrate external stimulation signals into each of the CTM populations by modifying **J**^**⋆**^***ϕ***^**⋆**^ in Eq. 11. By representing the stimulations applied to the cortical excitatory population, cortical inhibitory population, thalamic reticular nucleus, and thalamic relay nuclei as the firing rates *ϕ*_*x*_, *ϕ*_*y*_, *ϕ*_*z*_, and *ϕ*_*w*_, respectively, we can express the CTM equations as Eq. 15.

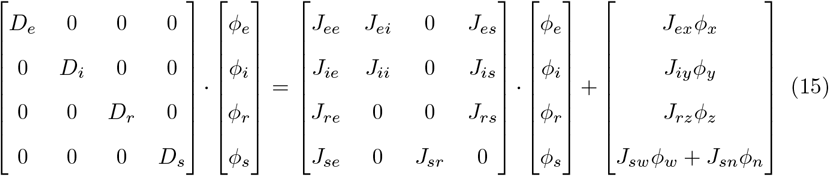

The solutions for the stimulation applied to each CTM population (separately) have a general form of:

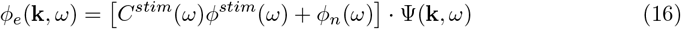

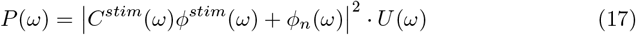

when *C*^*stim*^ is a stimulus-dependent function derived from Eq. 15. CTM power spectra for different stimulation targets are presented in Table 1.

**Table 1.**
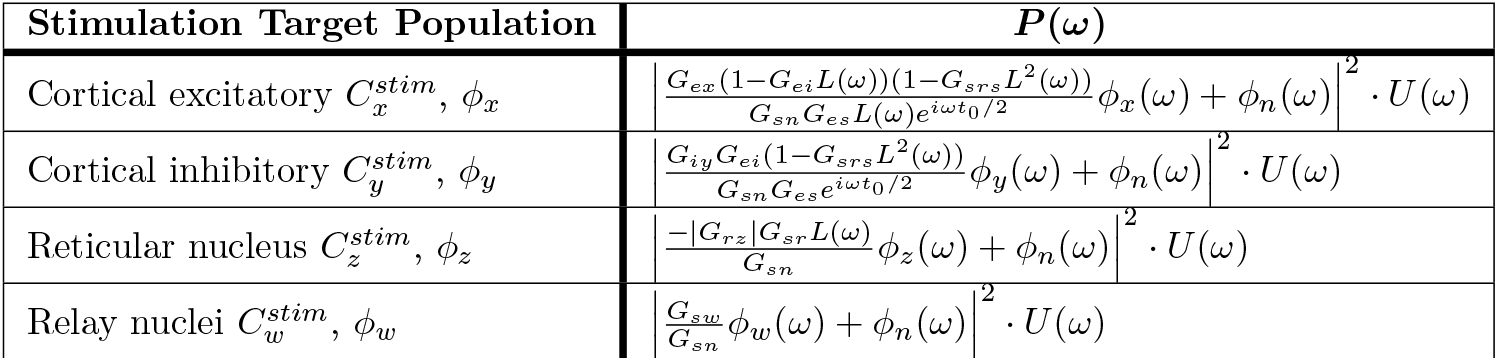
CTM power spectra for different stimulation targets.

In scenarios where a broad cortical region is stimulated, as is common in transcranial techniques such as rTMS or tACS, we assume that both the excitatory and inhibitory populations are excited concurrently by *ϕ*_*xy*_(*ω*). In this scenario, the power spectrum *P* (*ω*) can be expressed as follows:

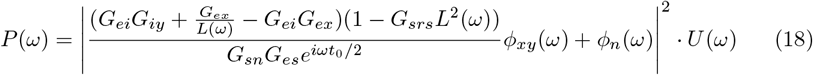

It is important to note that both rTMS and tACS typically target specific brain networks rather than the entire cortex, focusing on regions such as the dorsolateral prefrontal cortex that have been observed to have a greater impact on DoC patients compared to other areas [8, 18–20]. However, the CTM does not possess the multi-regional resolution required to simulate the cortex at this level of detail. Additionally, here, we do not consider potential variations in the response to TMS between inhibitory and excitatory populations. High-intensity stimulation primarily triggers firing in excitatory-to-excitatory axons due to their extensive spatial coverage and alignment with the electric field generated by TMS. In contrast, lower stimulation intensities are more likely to elicit firing events in inhibitory cells due to their lower firing thresholds [29].

### Stimulus equations

As previously mentioned, to derive the stimulus formula, we need to solve *P* ^*hlt*^(*ω*) = *P* ^*doc*+*stim*^(*ω*). Therefore, after fitting a CTM to a patient with DoC, we substitute *P* ^*doc*+*stim*^(*ω*) with Eq. 17, resulting in the following expression:

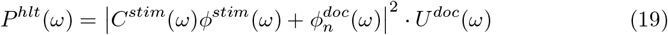

and

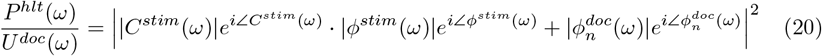

To extract |*ϕ*^*stim*^| in Eq. 20, we need to set |*ϕ*_*n*_| = 1, as previously mentioned. Additionally, we must determine the stimulus phase ∠*ϕ*^*stim*^(*ω*). Eq. 13 provides information about the amplitude at each frequency, but does not impose constraints on the phase. Therefore, we allow ourselves to set the phase in a manner that is instrumental in solving the equation above. Specifically, we set it as a function of 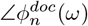 and add *π* to introduce a negative sign in *C*^*stim*^(*ω*), as shown in Eq. 21. For consistency with the theory, we set 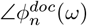 to be a random number with a uniform distribution since *ϕ*_*n*_ is approximated by white Gaussian noise.

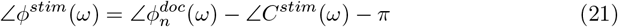

and get:

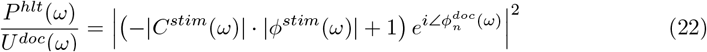

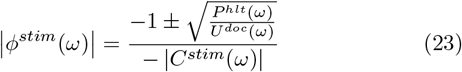

To obtain a positive power spectrum, we opted for a negative sign before the square root in the expression. Additionally, we can set 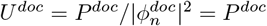, while keeping in mind that *P* ^*doc*^(*ω*) is an analytical formula. It is important to note that unlike *P*^*hlt*^ that can be either analytical or experimental, 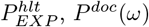 cannot be experimental since there must be consistency between the power spectra and *C*^*stim*^, which relies on fitted parameters.

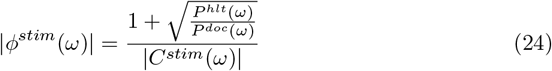

By performing an inverse Fourier transform of *ϕ*^*stim*^(*ω*), we construct the stimulus time series *ϕ*^*stim*^(*t*). Table 2 presents the amplitudes and the phases of stimuli applied to the various populations of the CTM.

**Table 2.**
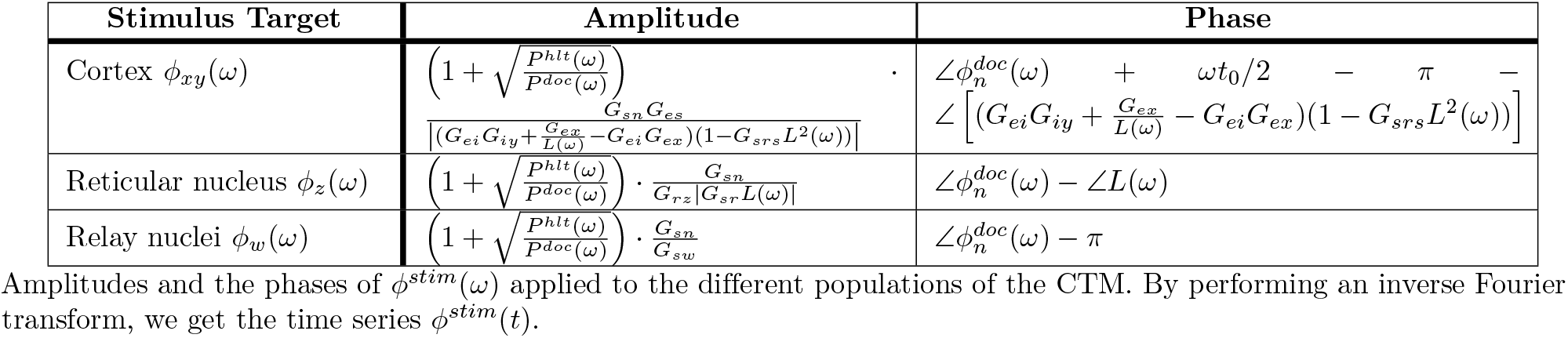
Stimuli amplitudes and phases.

At this point, we can also determine the stimulus intensity value for practical purposes, as outlined in the discussion section. This can be achieved by tuning *G*_*ex*_, *G*_*iy*_, *G*_*rz*_, and *G*_*sw*_, which were initially set to 1. However, it is worth noting that the value of *G*_*sn*_ was determined during the fitting process, as previously mentioned.

### EEG signal generation

We generated artificial EEG signals from “Healthy-fitted”, “DoC-fitted”, and “DoC-fitted with stimulation” models. These signals were generated for a duration of 30 seconds and were sampled at a rate of 125Hz. Since the generated signal achieves stationarity after 5 seconds (which were disregarded), this duration proves adequate for computing power spectra. The EEG signals were simulated on a 0.5 m *×* 0.5 m square sheet, covering 784 locations. To make the simulated signals comparable to the experimental data, we interpolated them to match the locations of the 137 experimental electrodes and high-pass filtered those above 0.2Hz.

### Transient and long-lasting changes

Further, we investigated whether the constructed stimulus signal could induce a long-lasting change in the model’s behavior, i.e., a bifurcation. We generated EEG signals from the “DoC-fitted” model for a duration of 150 seconds, while the stimulus was active during a specific time window from the 30th second to the 60th. We computed the spectrogram of the Cz electrode signal and examined the time periods before, during, and after the stimulation.

However, the model fitting process described previously ensures that a CTM fitted to steady-state activity remains stable and resistant to perturbations, making it challenging to induce changes solely through external stimulation. The stability of the CTM is determined by the denominator in Eq. 12. When all the zeros of Eq. 25 satisfy *Im*(*ω*_0_) *<* 0, the model is stable. Model stability boundaries can be represented in a three-dimensional space, as defined in Eq. 26. The *X,Y,Z* parameters correspond to corticocortical, corticothalamic, and intrathalamic loop strengths, respectively. These parameters offer a qualitative representation of the majority of CTM’s dynamics and provide insights into its stability characteristics across various parameter values [26]. By adjusting these parameters, we can move the fitted model beyond its stability region, allowing it to alter its state in response to stimulation.

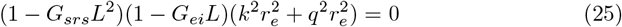

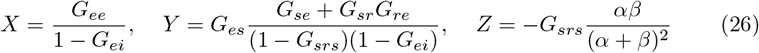

A notable stability region in the CTM is represented by the equation *X* + *Y <* 1 [52]. Therefore, to introduce instability and trigger a (saddle-node) bifurcation, we deliberately brought the model to a critical point *X* + *Y* = 1. This was achieved by setting the value of *G*_*ee*_ according to Eq. 27. However, since *G*_*ee*_ also affects the power spectra (see Eq. 12), this approach cannot be applied to a fitted model as it will cease to produce the desired spectrum unless the fitted model is already near a critical point (in which case modifying *G*_*ee*_ is not necessary to induce a bifurcation).

In summary, when *X* + *Y <* 1 the base model is too stable to sustain any changes induced by the stimulus. As an alternative path, we chose to modify the fitted parameters (i.e., the parameters that determine the DoC power spectrum, effectively changing it to the representation of a different, virtual patient) to illustrate that under the right conditions (i.e., *X* + *Y* = 1), our proposed method holds the potential to bring about lasting changes.

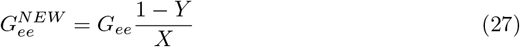

## Results

We successfully tested our methodology on subjects from the dataset. In the following results, we showcase its functionality using two pairs of DoC patients and healthy subjects. As evident from the calculations, the stimulus derivation procedure is nearly deterministic and can be applied to any pair of subjects with distinct power spectra. It is important to note that the non-deterministic component, particularly 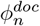, introduces some white noise to the output, but it does not alter the spectral shape [53].

### Power spectra

Fig. 5 illustrates the outcomes of stimulation applied to the thalamic reticular nucleus of the MCS patient and to the cortex of the UWS patient, as expressed in the power spectra. We constructed a stimulus time series for different regions of CTMs fitted to these patients. Upon applying the stimulation to the models, we observed that the power spectra of the simulated EEG closely resembled those of healthy subjects, exhibiting increased power at alpha and beta bands, characteristic of healthy individuals. Notably, all the stimuli constructed for the other CTM populations induced the same power spectra as the spectra presented in Fig. 5. Additionally, we generated stimulations based on an experimental healthy spectrum (see Fig. 5c). Although 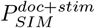 did not match 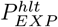 as closely as 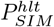, it still produced a satisfactory approximation. Overall, these outcomes highlight the efficacy of our approach in obtaining healthy-like EEG spectra in CTMs fitted to DoC patients.

**Fig 5.**
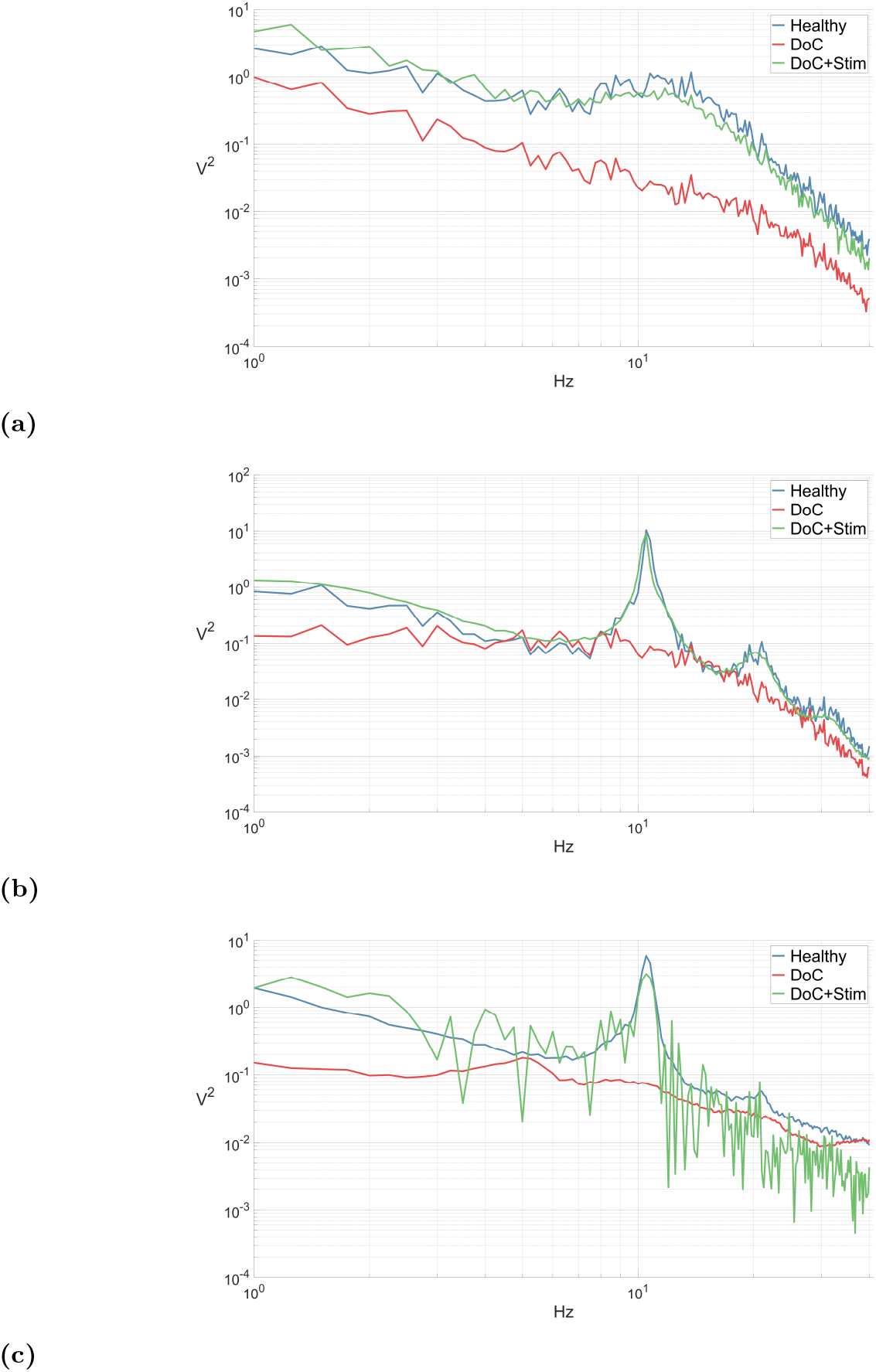
Power spectra comparison: examples of the stimulation on different patients and regions. The red curve represents the average power spectrum of a patient with DoC, the blue curve represents the average power spectrum of a healthy subject, and the green curve is the average power spectrum of a CTM-generated time series when the CTM is fitted to a patient with DoC and is stimulated in one of the CTM populations with the derived stimulus. **(a)** MCS patient and healthy subject #1: the stimulus is applied to the thalamic reticular nucleus. **(b)** UWS patient and healthy subject #2: the stimulus is applied to the cortex. **(c)** Same as *b*, but the red and blue curves are experimental EEG spectra, and 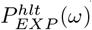) is used in the stimulus calculations.

### Phase constraints

In our experimental setup, the random number generator used for generating sensory inputs during model simulation was different from the one used for stimulus construction. This means that phase 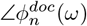 used in the injected stimulus was different from phase 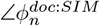 that fed the simulator, even though both followed a uniform distribution. This setup resembles a stimulation configuration when the real-time brain signal phases are not accounted for by the stimulation equipment. However, it contradicts our method’s specific requirement, which states that the stimulus phase ∠*ϕ*^*stim*^(*ω*) depends on the thalamic sensory input phase 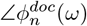 (see Eq. 21). Fulfilling this requirement is exceptionally challenging, especially *in vivo*, where measuring thalamic inputs in real-time poses significant difficulties. Yet, as long as the condition *P* ^*hlt*^(*ω*) ≥ *P* ^*doc*^(*ω*) (Eq. 19) is met for every *ω* (see Fig. 5a, for instance), this disparity does not significantly affect the results. To further prove this point we experimented with creating a stimulus with a linear phase instead of a random one, while keeping the sensory inputs as white noise. The stimulus time series appeared different (see Fig. 6a, 6b), but the resulting *P*^*doc+stim*^ in Fig. 5a remained unchanged.

**Fig 6.**
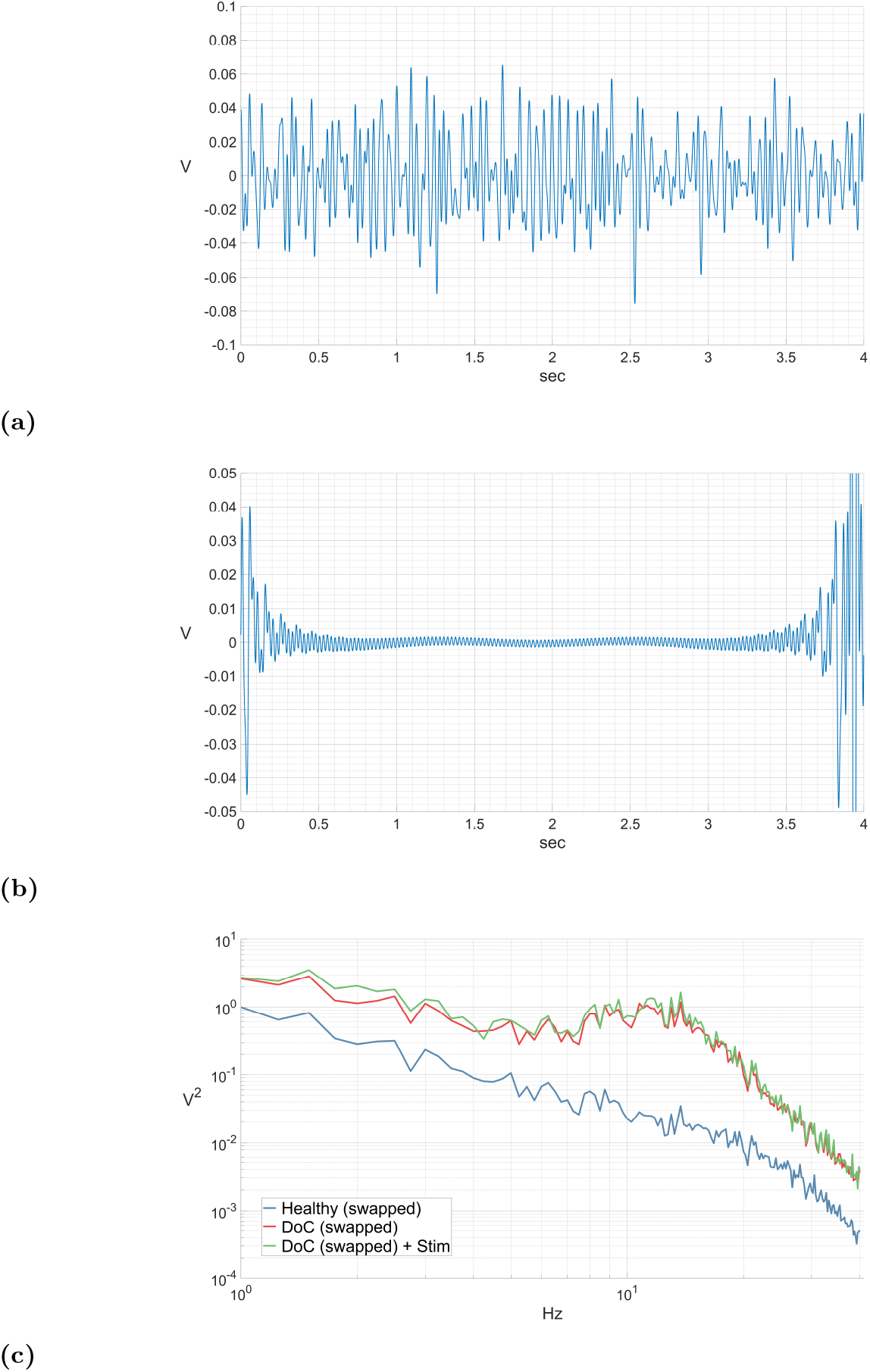
Stimulus phase effects. **(a) & (b)**: Time series of the stimulus to the reticular nucleus, derived for the MCS patient and healthy subject #1. The stimulus was calculated using 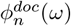 with a random phase **(a)** and a linear phase **(b)**. For all the different phases, the generated signal power spectrum appeared similar to Fig. 5a. In **(c)**, we swapped between healthy subject #1 and the MCS patient to demonstrate that for *P* ^*hlt*^(*ω*) *< P* ^*doc*^(*ω*), the stimulation fails due to a phase mismatch in the experimental setup.

However, when there are values of *ω* for which *P* ^*hlt*^(*ω*) *< P* ^*doc*^(*ω*), it becomes crucial to maintain the defined relationship between 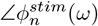 and 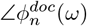. As shown in Eq. 19, *C*^*stim*^(*ω*)*ϕ*^*stim*^(*ω*) is added to the DoC power spectrum. It can either increase or decrease the power at different frequencies based on its phase; for example, adding a phase of *π* radians is equivalent to multiplying the amplitude by -1. In our experimental setup, since 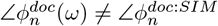, a power decrease does not occur at frequencies where *P* ^*hlt*^(*ω*) *< P* ^*doc*^(*ω*). An example illustrating this behavior can be seen in Fig. 6c, where we exchanged the roles of the healthy subject and the MCS patient (stimulating the healthy subject in the thalamic reticular nucleus) to create a scenario where this issue arises. In this case, *P*^*doc+stim*^ followed *P*^*doc*^ instead of *P*^*hlt*^, and the stimulation did not yield the desired effect.

### Hysteresis and bifurcating effects

Fig. 7a illustrates the changes in the CTM-generated signal spectrum before, during, and after the stimulation. The stimulus was administered to the thalamic relay nuclei in a model fitted to the USW patient from the 30th to the 60th second. Throughout the stimulation period, there was a substantial increase in power across all frequencies, reflecting *P* ^*hlt*^(*ω*) *> P* ^*doc*^(*ω*). Following the termination of the stimulus, the model exhibited a hysteresis effect that lasted for a 2-second transient period, presenting intermediate power values before returning to its pre-stimulation state. Hence, no long-term changes in the model’s output were observed following the stimulation.

**Fig 7.**
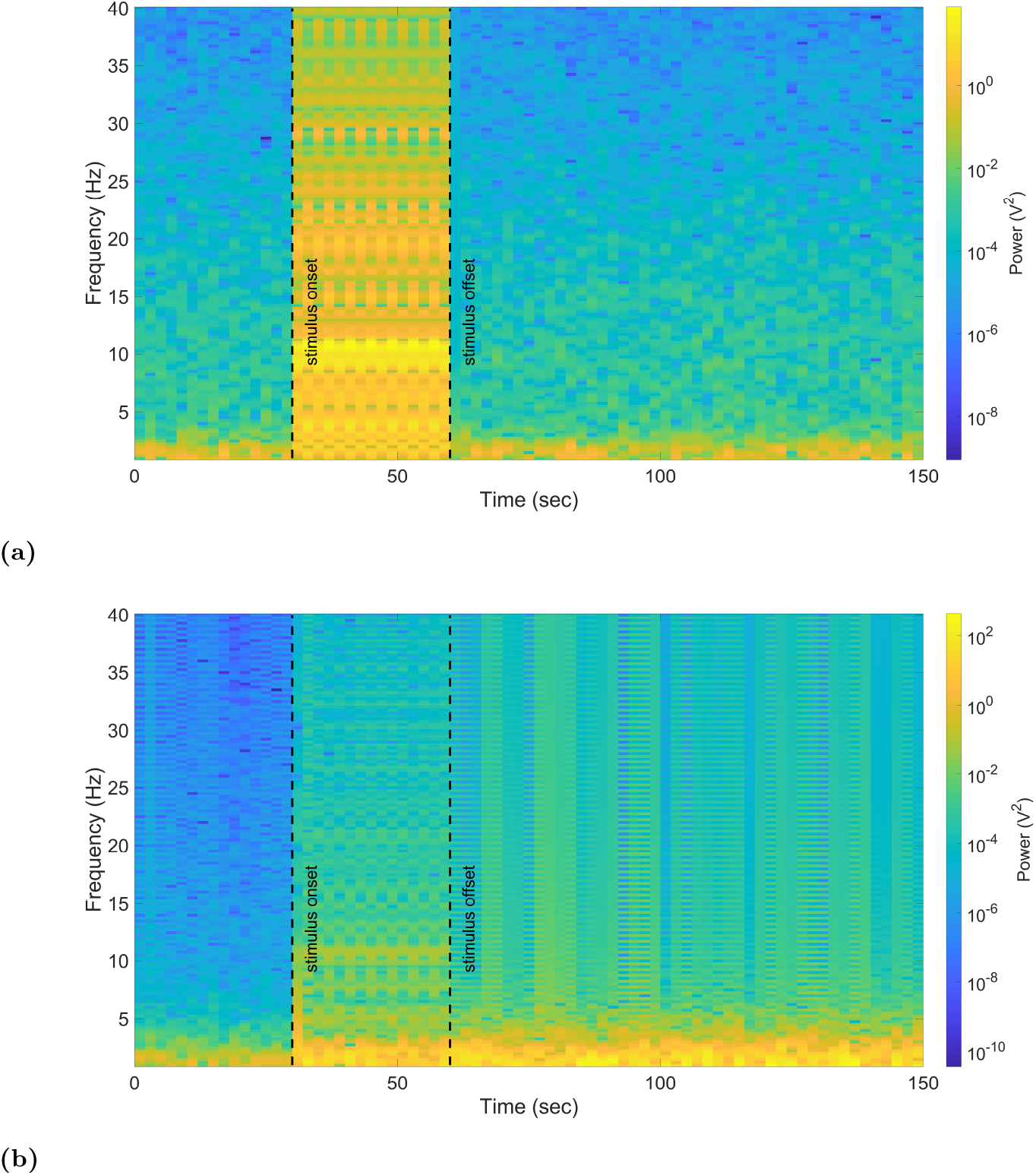
Transient and long-lasting changes. **(a)** A 150-second spectrogram of a Cz electrode EEG time series generated from a CTM fitted to the UWS patient. The model was stimulated in the thalamic relay nuclei from the 30th to the 60th second using a stimulus calculated based on 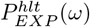 from healthy subject #2. **(b)** Same as, but the CTM was brought to a critical point before the stimulation.

However, when we deliberately brought the model to a critical point (*X* + *Y* = 1) where the CTM was less stable, a bifurcation occurred, and the model’s activity did not revert to its initial state upon termination of the stimulation, as shown in Fig. 7b.

During the stimulation period, the spectrogram looked similar to the non-critical case, but with a notable decrease in overall power, particularly above 4Hz. After the stimulus offset, the activity did not return to its initial state. Instead, it maintained power levels observed during stimulation, which were significantly higher compared to the pre-stimulation period. The pattern of the activity at the spectrogram appeared to combine the initial activity (or potentially slow-wave activity, given the CTM’s tendency towards it near *X* + *Y* = 1 [52]) with non-physiological artifacts.

## Discussion

Our study introduces a computational approach aimed at generating a signal capable of inducing healthy-like power spectra in patients with DoC. This involves fitting a CTM to the power spectrum of a patient with DoC and then utilizing the power spectrum of a healthy subject to derive a personalized stimulus time series. We applied our method to several patients with DoC, targeting brain regions commonly addressed in DBS, rTMS, and tACS. Consequently, these stimuli drove the CTM to generate EEG power spectra resembling those seen in healthy brains.

A central question arises: Will this method be effective *in vivo*? As highlighted in the introduction, various brain stimulation therapies have been employed in the treatment of DoC, but their success has been only partial [8]. Our stimulus construction technique, once integrated into these therapies, aims to enhance their outcomes by inducing brain activity in patients that closely resembles the patterns seen in healthy individuals. This is grounded in two key assumptions that warrant further examination:

- The proposed model has the capacity to capture the brain dynamics of DoC patients and healthy individuals. In other words, once we apply this stimulation method to a real brain, it will yield an activity akin to the one produced by the model. If not, we may need to consider employing a model that is more complex, as detailed below. However, this approach has its limits due to unknown empirical constraints that can’t be modeled and may prevent the brain from producing the desired activity.
- A healthy-like spectrum increases the likelihood of regaining consciousness. While this is both a philosophical and phenomenological query, the assumption is based on the research of neural correlates of consciousness [54–58]. Numerous studies have demonstrated significant differences in the power spectra of DoC patients compared to healthy controls [59–62]. While a state of consciousness frequently results in a power spectrum akin to that of a healthy individual, the opposite is not always true, and other factors might also be important (e.g., integration and segregation) [63]).

Furthermore, we propose that our personalized stimulation method, when used consistently over multiple sessions, may lead to longer-lasting effects compared to current stimulation approaches. We hypothesize that this method could stimulate the brain’s natural plasticity mechanisms by inducing activity patterns similar to those observed in a healthy brain. Such activation could potentially aid in the brain’s gradual recovery, helping it regain normal function without the continuous need for external intervention. [64, 65]. However, modeling such plasticity, as shown in prior studies [29, 66, 67], requires integrating time-dependent functions into the model parameters, and specifically involves applying concrete learning rules to adjust them.

Nonetheless, our results demonstrate that after the termination of the applied stimulus, the model’s activity reverts to its initial state, failing to undergo the necessary bifurcation required to sustain healthy-like activity. Conversely, when we brought the model to a critical point prior to stimulation, a bifurcation occurred, resulting in lasting and substantial changes in the generated signal. Although these changes were significantly different from the desired healthy-like activity, and bringing a fitted model to a critical point is incompatible with our method, this experiment underscores the model’s capacity to undergo bifurcations in response to the constructed stimulus. It is possible that a more complex model could exhibit bifurcations that lead to the desired sustained activity when subjected to the constructed stimulation [68]. However, a comprehensive analysis of CTM dynamics is necessary to better understand the stimulation effects, as manifested by the trajectories within the *XY Z* parameter space (see Eq. 26) [25, 69]. Given all that, it is important to clarify that the limitation of the current model to present long-lasting effects does not negate the potential for such effects in our approach *in vivo*.

### Increasing model complexity

As previously mentioned, it may be necessary to use a more intricate model to refine the stimulation and control its effects, thus improving the chances of success of this method in an *in-vivo* setting. The CTM, while capable of replicating various brain states and phenomena effectively, is inherently simplistic [24]. A more sophisticated model has the potential to offer a more accurate representation of both DoC and healthy brains in terms of the desired bifurcations and neural plasticity. For instance, whereas the CTM is adequate for modeling typical sleep stages and wakefulness, unconventional arousal conditions like jet lag or caffeine intake necessitate the incorporation of additional dynamics and bifurcations. This can be achieved by establishing connections between the brainstem and hypothalamus with the CTM to drive transitions between different states, as demonstrated by Phillips et al. [41, 68].

Furthermore, integrating additional brain regions into the CTM and enhancing the model resolution by subdividing its components would expand the scope of modeled stimulation techniques, facilitate targeted stimulation modeling and potentially streamline the application of such stimulations more effectively. Specifically, incorporating the vagus nerve and the brainstem into the CTM would enable support for VNS. Segmenting the cortex into multiple populations would aid in modeling focused stimulation of specific brain networks as commonly practiced in rTMS and tACS. With detailed network modeling, which may encompass multiple cortical areas, various nuclei and ganglia (as in the case of a stimulation model for Parkinson’s disease [31]), the most effective stimulation sites can be accurately identified, thereby guiding the placement of electrodes for a targeted DBS, or instead opting to apply a wide-angle rTMS [8, 9, 18]. Additionally, it allows for the simultaneous stimulation of different brain areas and for exploration of how stimuli propagate through the brain.

Another important motivation is related to the etiology of the DoC state. DoC can arise from diverse causes such as traumatic brain injury, anoxia/hypoxia, ischemic/hemorrhagic stroke, intoxication, and more [1]. Consequently, each DoC patient may present with a distinct etiology characterized by brain lesions, disrupted structural and functional connectivity, spatial variations in the cortex, and other factors. Given that stimulation is typically applied to healthy brain regions [70], modeling these specific pathologies is essential to tailor patient-specific stimulation.

Finally, to integrate our stimulus-derivation method into brain stimulation therapies like DBS or tACS, we must provide a model for them. This is necessary because these therapies involve constructing and transmitting stimuli in the form of electric currents or magnetic fields, whereas in our method the applied stimulus is represented as a firing rate *ϕ*^*stim*^(*t*). The model needs to encompass parameters for constructing the signal for different stimulation protocols, modulatory effects of the mediums it propagates through, its influence on the neural activity of the target population (i.e., its translation to *ϕ*^*stim*^(*t*)), and additional side effects like tissue heating that may impact subsequent stimuli signals. Constructing such a model is possible within the NFT framework, as partially demonstrated in previous studies for DBS and rTMS [29, 31]. Such a model can also be created for non-electromagnetic therapies like low-intensity focused ultrasound pulsation, recently used as a therapy for DoC [6, 7, 71].

### Practical adaptations for *in-vivo* application

#### Modeling and tuning brain stimulation therapies

We plan to incorporate our stimulation approach into DBS, rTMS, and tACS brain stimulation therapies, however, as mentioned above, we first need to develop models for them. Then, we can invert these models to determine the stimulation protocols and their parameters based on *ϕ*^*stim*^(*t*). The precise mechanisms underlying the effects of these stimulation modalities on neurons are still under investigation [17, 72, 73].

However, we can broadly model DBS and rTMS as spike-like signals with a firing rate of *Fr*(*t*) = *ϕ*^*stim*^(*t*) and tACS as an electric potential *V* (*t*) = *ϕ*^*stim*^(*t*). Consequently, in DBS and rTMS, *ϕ*^*stim*^ modulates the stimulation frequency (akin to an FM radio) through the alteration of inter-pulse intervals, whereas in tACS, it directly determines the signal.

The configuration of the stimulation parameters should be based on both *ϕ*^*stim*^(*t*) and clinical requirements. Fine-tuning the stimulation power and frequency to align with individual patient needs and specific medical conditions is a standard practice in brain stimulation therapies, aimed at optimizing treatment outcomes, while minimizing adverse effects [74–77]. In the proposed models, the chosen stimulation frequency in DBS and rTMS (the “carrier” frequency), and the direct current level in tACS correspond to the average intensity of *ϕ*^*stim*^, regulated by *G*_*ex*_, *G*_*iy*_, *G*_*rz*_, and *G*_*sw*_.

However, as outlined in the methods section, the stimulus intensity is predetermined and should be set according to therapy properties, rather than vice versa. Therefore, we utilize the tuned stimulation power and frequency to determine the values of *G*_*ex*_, *G*_*iy*_, *G*_*rz*_, and *G*_*sw*_.

#### Sensory input phase acquisition

As outlined in Eq. 21, the thalamic sensory input phase 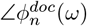 is necessary for the calculation of *ϕ*^*stim*^. We demonstrated that when *P* ^*hlt*^(*ω*) ≥ *P* ^*doc*^(*ω*), it is possible to set any arbitrary phase instead of 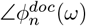 during the stimulus calculation. However, when this condition is not met, precise knowledge of the sensory input phase becomes crucial.

Theoretically, acquiring the sensory input phase would require measuring the activity of all the thalamic afferent neurons, which is not feasible with present technology. Nonetheless, strides have been made in this direction, such as estimating the phase from population activity and synchronizing DBS signals with neural activity for more efficient Parkinson’s disease treatment [78]. Alternatively, selecting a high-power *P*^*hlt*^ could circumvent this obstacle.

#### Closed-loop stimulation

Our method can be combined with closed-loop stimulation, a paradigm for personalizing neuromodulation that has made a lot of progress in recent years. Grounded in control theory, closed-loop brain stimulation involves the continuous monitoring of brain activity, providing real-time feedback to adjust stimulation parameters. This approach has significantly improved therapies like DBS, rTMS, and tDCS, enhancing response rates and reducing outcome variability for conditions such as Parkinson’s disease, epilepsy, depression, and DoC [71, 79–81].

In our method, real-time closed-loop feedback can maintain the patient’s power spectrum at the desired level during stimulation. Any deviations will dynamically adjust the intensity of *ϕ*^*stim*^ through the parameters *G*_*ex*_, *G*_*iy*_, *G*_*rz*_, and *G*_*sw*_. These adjustments, in turn, will regulate the “carrier” frequency or the direct current level, depending on the specific stimulation therapy used.

In order to fully adapt the stimulus for real-time closed-loop feedback, the model parameters need to exhibit temporal variability, i.e., representing plasticity effects. Without this adaptation, our current stimulus construction method does not allow parameters to be updated in real-time during stimulation, as the available feedback biomarker is the resting-state EEG spectrum, obtained when stimulation is off, possible only between therapy sessions. Temporal variability in parameters will transform the power spectrum into a dynamic equation that integrates feedback inputs. This also suggests that real-time closed-loop feedback will enhance the performance of models incorporating plasticity compared to periodic updates every few minutes.

### Further applications and extensions

As we previously mentioned, our proposed approach offers versatility and can be adapted to various spectra and applied to different brain regions modeled by NFT. It is evident from the calculations of the stimulus derivation, that they can be applied to any two individuals, regardless of their medical condition. Essentially, with the EEG power spectra of two individuals, we can construct a stimulus signal that will cause person #1 to induce the power spectrum of person #2, for as long as the stimulus is applied. The only constraint is imposed by the availability of a sensory input phase, required for cases when *P* ^#2^(*ω*) *< P* ^#1^(*ω*). Moreover, NFT models can be fitted to power spectra acquired through other neuroimaging techniques such as magnetoencephalography and intracranial electroencephalography, in addition to electroencephalography [24]. Therefore, the potential application of our method extends beyond DoC, making it suitable for addressing a broad range of medical conditions across diverse imaging and stimulation settings.

Furthermore, as stated above, the model can be expanded by incorporating additional brain regions and stimulating them as desired [31]. For instance, we can create personalized stimuli for DBS treatment for depression, where remission rates are just above 50%. This extension would involve modeling the limbic system and stimulating regions like the striatum and subgenual cingulate cortex. Although the equations will differ, the core approach remains consistent, with the use of *P*^*depression*^ in place of *P*^*doc*^. Our method has significant potential to improve treatment outcomes by enhancing the longevity of therapeutic effects and aiding in target selection [82, 83].

Finally, it would be intriguing to investigate whether, by toggling between *P*^*hlt*^ and *P*^*doc*^ (and accounting for sensory input phase 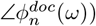, one could induce a reversible anesthesia-like state in a healthy subject. This exploration could shed light on the potential for this method to modulate brain activity across various physiological and altered states.

## Summary and conclusion

In this research, we introduced a novel computational method aimed at modulating brain activity in patients with DoC. Our approach focused on constructing a stimulus signal that could induce a healthy-like neural activity pattern in these patients. To achieve this, we utilized a simplified brain model based on NFT and adapted it to reproduce the power spectrum of DoC patients. The core idea of our method involved fitting this NFT model to the power spectrum of the patients and, subsequently, using the power spectrum of healthy subjects to derive stimulus time series. By applying the derived signal to brain regions typically targeted by therapies like DBS, rTMS, and tACS, we demonstrated that our stimuli induced EEG power spectra resembling those of healthy individuals. We speculate that once the brain produces a healthy-like activity, plasticity takes place, which is likely to cause long-lasting changes and improvement in a patient’s consciousness level compared to current stimulation approaches.

While our results provide promising insights into the potential of this approach, several critical considerations and avenues for future exploration should be noted. Firstly, the development of various stimulation therapy models needs to be pursued and integrated with the proposed NFT model to bridge the gap between theoretical frameworks and practical applications. Moreover, the implementation of our method in clinical settings requires further validation and a thorough comparison with the outcomes of existing stimulation protocols to ensure its efficacy and safety. Key questions include whether the computational model accurately represents the complexity of the DoC brain and whether the assumption that a healthy-like power spectrum implies consciousness holds true *in vivo*.

Additionally, we discussed the importance of testing more complex models to capture the intricate nature of DoC and to enable targeted stimulation. Detailed models could help determine optimal stimulation locations and patterns, thereby enhancing the efficacy of brain stimulation therapies. Furthermore, understanding the specific etiology of DoC in an individual patient and modeling those associated pathologies may be crucial for personalized stimulation strategies.

It is important to emphasize the generalizability of our stimulus derivation technique, as its computations are deterministic and adaptable to any pair of subjects with different power spectra. The only constraint in certain cases is the availability of sensory phase information. Therefore, this technique can be tailored to address other pathologies treated with brain stimulation beyond DoC, opening up a multitude of potential avenues for exploration.

In conclusion, our research provides a proof-of-concept computational framework for designing patient-specific brain stimulation therapies. While further research specifically aiming at the clinical translation is needed, this method holds promise for improving the outcomes of future neuromodulation therapies for DoC patients and could be extended to treat other conditions amenable to brain stimulation interventions.

## Acknowledgments

We would like to thank Avigail Makbili from the Computational Psychiatry Lab at Ben-Gurion University, who took part in the processing and cleaning of the experimental EEG data. Additionally, we want to thank Prof. Yaniv Zigel from the Department of Biomedical Engineering at Ben-Gurion University, who helped develop the mathematics of this method.

Funding for this work was provided by a grant from the EU CHIST-ERA program. Additionally, this work was supported by the University and University Hospital of Liège, the Belgian National Funds for Scientific Research (FRS-FNRS), the FNRS PDR project (T.0134.21), the ERA-Net FLAG-ERA JTC2021 project ModelDXConsciousness (the Human Brain Project Partnering Project), and the FNRS MIS project (F.4521.23), JA is postdoctoral fellow funded (1265522N) by the Fund for Scientific Research-Flanders (FWO), O.G. is a research associate at FRS-FNRS.

The wording revision process of this article included the use of ChatGPT3.5 by OpenAI.

